# MINSTED tracking of single biomolecules

**DOI:** 10.1101/2023.09.15.557902

**Authors:** Lukas Scheiderer, Henrik von der Emde, Mira Hesselink, Michael Weber, Stefan W. Hell

## Abstract

We show that MINSTED localization, a method whereby the position of a fluorophore is identified with precisely controlled beams of a STED microscope, tracks fluorophores and hence labeled biomolecules with nanometer/millisecond spatio-temporal precision. By updating the position for each detected photon, MINSTED recognizes fluorophore steps of 16 nm within < 250 microseconds using about 13 photons. The power of MINSTED tracking is demonstrated by quantifying side-steps of the motor protein kinesin-1 walking on microtubules and switching protofilaments.

---

Measuring conformational changes and movements of individual proteins and other biomolecules is key to understanding their function. A powerful approach to this end is labeling the biomolecule with a fluorophore and tracking its position with an optical microscope. In contrast to scattering gold or latex beads which generate much higher photon rates, fluorophores are much smaller than proteins and can be specifically linked to numerous protein sites with minimal functional interference. On the other hand, fluorophores bleach and entail fluorescence intermissions. Moreover, they produce relatively low photon rates, which is disadvantageous because established methods for optical localization require a large number of detected photons *N* from a given position. This applies to both the popular localization by centroid calculation of the fluorescence diffraction pattern on a camera^1-3^ and the alternative method of scanning a Gaussian excitation beam around the fluorophore in a confocal microscope^4^. The need for large *N* is exacerbated by the fact that in both methods the localization precision scales with 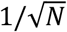. Doubling the precision therefore requires four times more detected photons, which usually entails four times longer periods of uninterrupted fluorescence and a temporal resolution that is deteriorated accordingly.

A way out of this catch is offered by localization through optical coordinate-targeting as realized in the methods called MINFLUX^5,6^ and MINSTED^7^. These methods harness a laser in order to optically establish and target a reference coordinate in the focal plane with nearly infinite precision. For this purpose, a laser beam is modified usually such that it forms a donut with a central point of minimum (ideally zero) intensity in the focal plane. Serving as a reference point, the minimum is rapidly steered with Angström accuracy around the fluorophore. Localization then boils down to finding out the unknown position of the fluorophore relative to the known position of the donut minimum. This relative position is readily assessed from the fluorescence rate, because this rate is determined by the well-known intensity gradient around the donut center. Ideally, coordinate-targeted localization is performed iteratively, by continually relocating the donut minimum such that the average distance between the minimum and the fluorophore becomes smaller in each iteration. At the same time, the donut intensity is increased so that the fluorescence rate stays largely constant. As a result, the localization precision scales with *e*^−*N*^ (ref.^8,9^), rather than slowly with 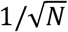, making coordinate-targeted localization with a given *N* more photon-efficient.

In MINFLUX, the wavelength of the laser donut-beam excites fluorophores. Therefore, the fluorophore is at the donut position where its fluorescence rate would be minimal. In contrast, MINSTED uses a regularly focused beam for excitation and the donut is used for de-excitation through stimulated emission depletion (STED)^10^. Hence, in MINSTED the fluorophore is at the donut position where the fluorescence rate would be maximal. Yet despite this difference, both MINFLUX and MINSTED attained Angström localization precision in superresolution imaging^5,11^. In addition, taking advantage of the fact that current MINFLUX implementations need about 100 times fewer fluorescence photons than established localization methods, MINFLUX was shown to track labeled proteins with record (sub)millisecond per nanometer spatio-temporal precision^9,12^. This accomplishment brings up the question whether and to which extent MINSTED is suitable for molecular tracking, given that its point of reference is a fluorescence maximum rather than a minimum. Here we show that owing to the virtues of coordinate-targeting, MINSTED attains a similar spatio-temporal resolution as MINFLUX. Moreover, we show that updating the position estimate of the emitter with each detected photon provides an elegant way of following the molecule’s motion directly, without additional computation. The power of MINSTED tracking is highlighted by revealing rarely observed leaps of the motor protein kinesin-1 when walking on microtubules, such as sudden protofilament switching and side-stepping.

## Results and Discussion

Like in STED microscopy, which has also been used to monitor one-dimensional protein movements by repetitive line-scan acquisition^13^, in single fluorophore localization by MINSTED the donut-shaped STED-beam reduces the space from which detected photons originate to sub-diffraction dimensions. This space is described by the so-called effective point spread function (E-PSF, Fig. 1a), giving the normalized probability of the fluorophore to emit a photon in that area. Scouting for the fluorophore position with the tightly co-aligned excitation and STED-beams is then tantamount to probing the position with the sub-diffraction-sized E-PSF. The emitted photons carry more positional information than in conventional localization approaches, with the improvement effected by continually reducing the full-width-half-maximum (FWHM) of the E-PSF through increasing the power of the donut-beam. At the same time, the donut minimum is moved closer to the fluorophore so that the emission rate of the fluorophore largely remains the same. Note that an added benefit of STED is that the donut keeps the (fluorescence) background low due to its innate signal-suppressing capabilities, which may bring about significant advantages over MINFLUX in many applications.

**Fig. 1.**
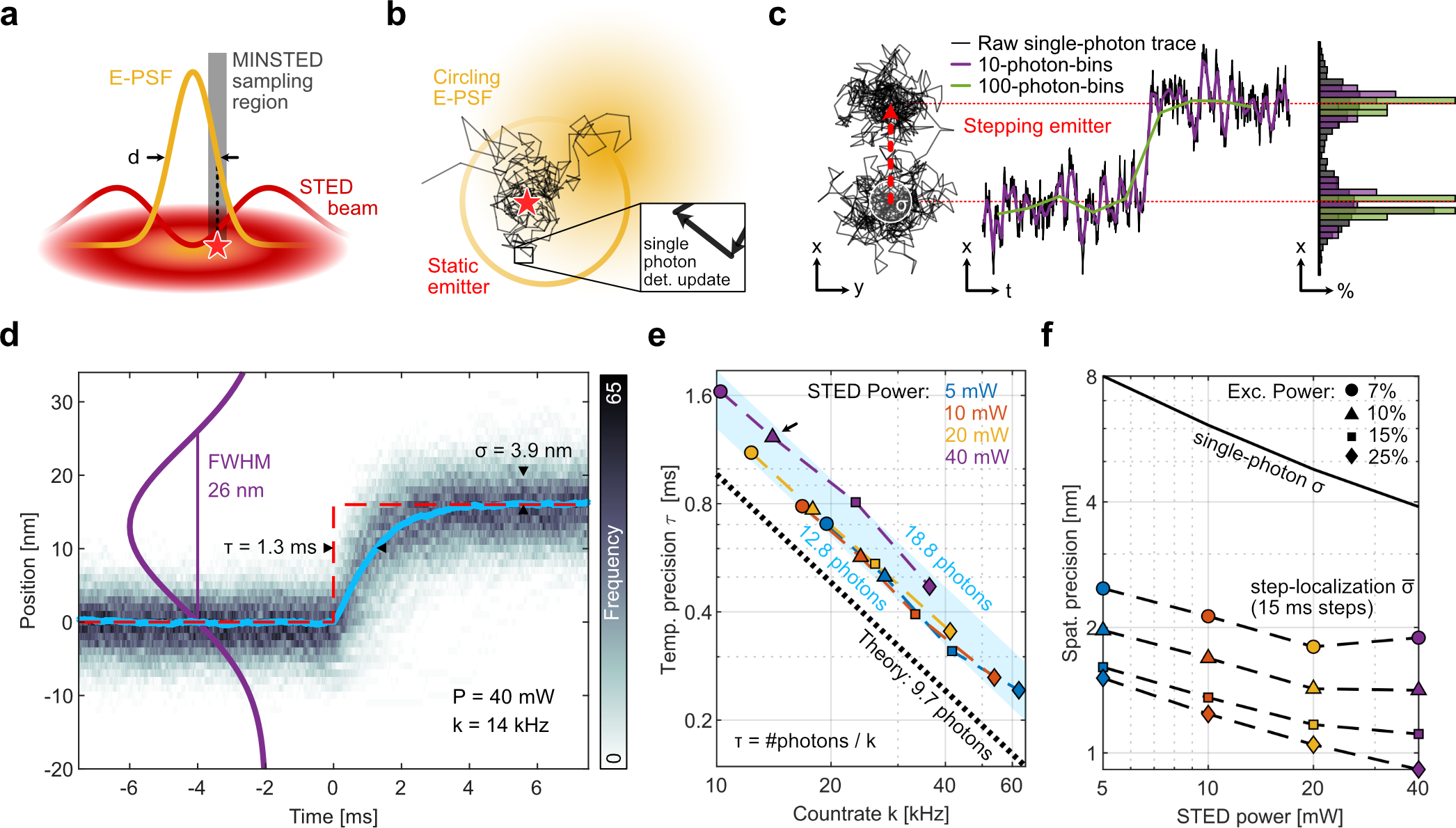
MINSTED localization principle and spatio-temporal precision. **a**, The STED E-PSF (orange) in the focal plane of the objective lens describes the probability that a fluorophore (red star) located in the region around the center of the donut-shaped STED-beam (red) emits a fluorescence photon. The E-PSF results from the joint action of a regularly focused beam for fluorophore excitation with a tightly co-aligned STED beam of longer wavelength for de-excitation. If covered by the donut crest (dark red) the fluorophore is essentially non-fluorescent, whereas at the central intensity minimum of the donut the fluorophore can fluoresce normally. In the MINSTED localization process, the co-aligned beams and thus the fluorescence probability given by the E-PSF, are ideally placed so that the fluorophore experiences the steep edge of the E-PSF (grey sampling region). **b**, MINSTED localization is performed by scanning the co-aligned beams, and thus the E-PSF (yellow), in circles around the assumed fluorophore position (star), such that the fluorophore keeps experiencing the E-PSF edge. The circle center is regarded as the (interim) localization. Upon detection of a photon, the circle center is shifted towards the latest scanning position so that it comes closer to the fluorophore. Thus, in a localization trace each ‘line’ indicates the detection of a photon and the ensuing change in center position (inset). Concomitantly, the STED-beam intensity is slightly increased so that the E-PSF becomes narrower and the circle diameter smaller, until a preset minimal FWHM value *d* is reached. With any additional detection, the circle center converges on the fluorophore position with a precision α. **c**, MINSTED tracks a sudden jump in position (step) of a fluorophore with a few photon detections. Binning the circle centers (rendered for each detection) increases the visibility of the step while compromising temporal information (purple and green lines). Data in **a-c** are based on simulations using setup parameters. **d**, Measured response to a quasi-instantaneous 16 nm step (red line) emulated by instantly displacing the co-aligned beams from a position matching the fluorophore and monitoring the relaxation of the beams towards the fluorophore by MINSTED localization. Overlaying 389 traces (histogram frequency shown as grey scale) at a STED beam power *P* = 40 mW, E-PSF FWHM of 26 nm, and a fluorescence countrate of *k* = 14 kHz reveals a nearly exponential average response function (blue) with a τ = 1.3 ms decay. The single-photon spatial precision α = 3.9 nm provided by MINSTED is estimated from the standard deviation of center positions along each trace after convergence. **e**, Temporal precisions τ as a function of the photon count rate *k* for varying STED power (color-coded), extracted from 16 sets of traces at different STED and excitation beam powers; *τ* scales with k^-1^; the arrow indicates the data point representing the data of panel d. The blue shading indicates the span of minimal (12.8) and maximal (18.8) average number of photons needed for convergence to the new position (defined by 1/e). The ideal *τ* (theory) is shown as black dashed line. **f**, Single-photon precision and step-localization precision as a function of STED beam power with excitation beam power as parameter. The single-photon-precision αand the step localization precision α scale with the STED beam power defining the FWHM of the E-PSF. α also scales with the number of photons detected per inter-step plateau and thus also with the excitation beam power indicated as percentage of 30 μW. Detecting < 500 photons suffices to localize 16 nm step plateaus with 0.8 nm precision.

In principle, the information carried by a detected photon is maximal when the FWHM is down to fluorophore dimensions. However, as the information from all the photons detected in the process contributes to the localization, molecular precisions are reached much earlier in the MINSTED localization process^11^; in fact, downsizing the FWHM to 26 nm suffices to reach 1 nm localization precision with *N* < 500 detected photons. In our implementation, the E-PSF is scanned on a circular trajectory around the assumed fluorophore position so that upon each photon detection, the position of the circle center is slightly shifted towards the latest scanning position (Fig. 1b). This procedure ultimately aligns the (average) circle center with the fluorophore position with continually increasing precision. Every circle center thus represents a position estimate based on the prior position and the latest photon detection. Once the E-PSF is down-sized to a minimal FWHM *d*, each detected photon gives an update of the fluorophore position with sub-diffraction precision. This single-photon-based localization update renders our MINSTED implementation particularly suitable for tracking (Fig. 1b).

Akin to any other optical localization approach, the temporal resolution in MINSTED tracking scales inversely with the photon detection rate *k*. Likewise, by averaging over consecutive detections the spatial precision can be improved at the expense of temporal precision. Yet, a key indicator of performance is the step response, quantifying how rapidly a localization method can measure an instant fluorophore displacement (Fig. 1c). To measure the step response, we employed the fluorophore Cy3B on a MINSTED setup featuring beam wavelengths of 560 nm and 636 nm for excitation and STED, respectively. Assuming a standard deviation α_*E*_ = FWHM/2.35 of the E-PSF and a sampling radius *r*, the response to an instantaneous displacement *s* ≪ *r* in the focal plane can be described by an exponential function with a temporal ‘decay’ constant 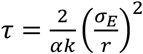. Here, α denotes the fraction of the scan circle radius *r* by which the circle is displaced after each detection. The radius *r* is proportional to α_*E*_, with *r/*FWHM = 0.5, meaning that τ scales with *k*^−1^ irrespective of the FWHM of the E-PSF. Rather than dislocating the fluorophore itself, which is technically difficult to execute sufficiently fast, we dislocated the co-aligned beams from a stationary fluorophore by 16 nm. This dislocation took only about a microsecond because our setup employed electro-optic beam deflectors. The subsequent localization trace, describing the (sub)millisecond convergence of the beams towards the original position, revealed the step response (Fig. 1d).

Averaging over multiple (389) responses, taken every 15 ms, validated the expected exponential behavior. Moreover, recording the responses for four different excitation and STED powers allowed us to extract τ (Fig. 1e). The evaluation showed that τ generally behaved as predicted. At the largest STED-beam average power (*P* = 40 mW, FWHM = 26 nm; *r* = 13 nm), however, τ tended towards higher values, accounting for the fact that the 16 nm step size is non-negligible with respect to the FWHM and the scan radius. Since *ss* ≪ *r* is no longer valid, the response time is not only limited by *k*, but also by the number of relocations of size αα*r* that the co-aligned beams need to take in order to cover the distance *ss*. Therefore, for some applications, a somewhat larger FWHM is more favorable (Fig. 1e). In any case, at *k* = 62 kHz, MINSTED detects 16 nm steps with τ < 250 μs.

The single-photon spatial precision σ is calculated as the standard deviation of the measured steps from the mean step response. It is independent of the detection rate *k*, which scales with the excitation power. The localization precision α of the coordinate of the step itself, which is obtained by averaging over the inter-step duration, scales with the number of the detections at a given step coordinate and hence with *k*^−0.5^. Averaging over a step of 15 ms yields 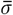 nm for *k* = 36 kHz and *P* = 40 mW. For a fluorophore that carries out steps *ss* ⪆ 2α, the steps can be directly identified in the raw data (or after binning over about 10 detections). Therefore, when measuring discrete fluorophore steps with s ⪆ 2σ, both the temporal precision (τ), as well as the inter-step averaged spatial precision 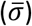, which depends on the duration of the current step, can be gained simultaneously. While the stepping times can be directly extracted from the raw single-photon trace, the averaging can be performed afterwards.

To explore the power of MINSTED for protein tracking, we resolved the steps of the motor protein kinesin-1 walking on microtubules. For this, we used a polymer-based in vitro assay consisting of an oxygen plasma-cleaned coverslip coated with the co-polymer PLL-PEG-biotin. Biotinylated and fluorescently labeled microtubules were immobilized to the surface via neutravidin; the surface between microtubules was blocked with biotinylated BSA (BSA-bt). The measuring buffer contained fluorophore-tagged kinesin-1, dithiothreitol (DTT) to reduce disulfide bonds, paclitaxel (Txl) to stabilize the microtubules, BSA-bt to reduce unspecific binding and a physiological (1 mM) concentration of adenosine-5’-triphosphate (ATP). To minimize fluorophore bleaching, oxygen was depleted by adding an oxygen scavenger system (OSS) consisting of pyranose oxidase, catalase and glucose. Adding a reducing and oxidizing system (ROXS) consisting of ascorbic acid (AA) and methyl viologen (MV) precluded the population of the fluorophore triplet state and enhanced the fluorescence rate.

Our tracking experiments (Fig. 2) were performed with three different constructs of kinesin-1 labeled with a single fluorophore at different sites via maleimide coupling: in the front (E215C), in the rear (K28C) and in the middle (T324C) of the head, with respect to the motor’s walking direction. As fluorophores, we used either Cy3B or ATTO 647N. For ATTO 647N, the excitation and STED wavelengths were 635 nm and 775 nm, respectively.

**Fig. 2.**
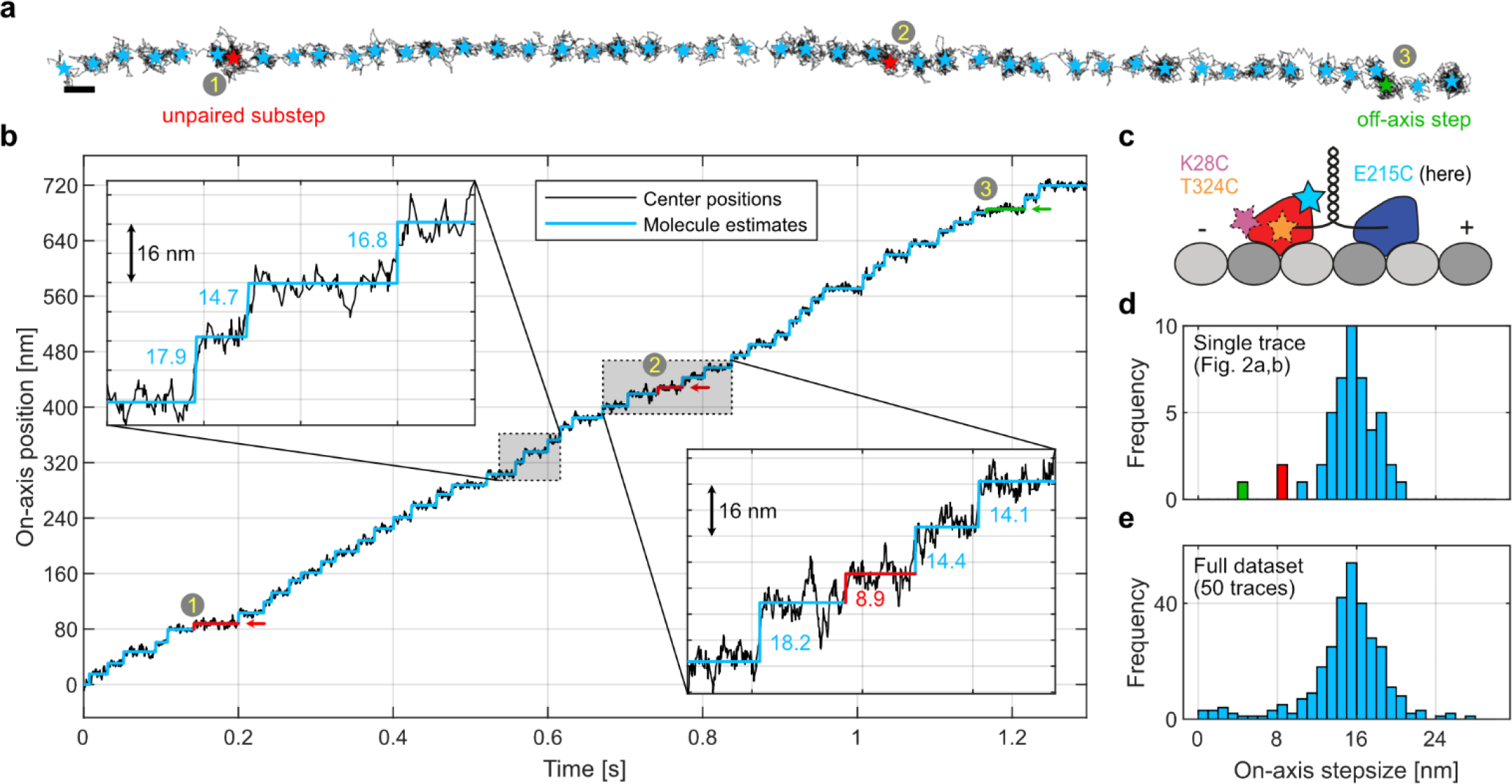
MINSTED tracking of kinesin-1. **a**, Stepping movement of kinesin-1 on a microtubule in the focal plane, with a single head labeled. Localizations (i.e. circle center positions) are shown in black; light-blue stars indicate inter-step mean positions. The numbers 1 and 2 (red stars) indicate a step size of ≈ 8 nm along the microtubular axis (i.e. on-axis); they can be attributed to kinesin reversing the order of its labelled and non-labelled head. Number 3 (green star) indicates a step that is mainly off-axis. Scale bar 16 nm. **b**, On-axis position of circle centers (black) along with their inter-step average (light blue) positions. The numbers refer to the same steps as in **a**, while the insets show zoom-ins with step sizes given in nm. **c**, Sketch of kinesin-1 homodimer and fluorophore position depicted as colored star in a given kinesin-1 construct; here, the E215C construct and the fluorophore ATTO 647N are used. **d**, Histogram of the on-axis step sizes of the displayed trace. **e**, Histogram showing on-axis sizes of steps from all MINSTED traces of E215C-ATTO 647N recorded.

With E215C-ATTO 647N, we recorded traces up to 736 nm in length with a temporal resolution of ∼1 ms, displaying clearly identifiable steps (Fig. 2a,b); see Methods for step detection. The step size histogram (Fig. 2d) of the shown trace displays a pronounced peak at around 16 nm, corresponding to twice the distance between the successive binding sites of a kinesin head on the microtubule. This observation is in good agreement with previous findings of the kinesin heads usually taking 16 nm steps at high ATP concentrations ^9^. Yet, the shown trace also displays two ∼ 8 nm steps at the 80 nm and 416 nm position coordinate on the microtubule (Fig. 2a,b (1,2) marked in red). As they are preceded and succeeded by regular 16 nm steps, we interpret these steps to arise either from the motor switching to a neighboring protofilament or from transiently detaching from the microtubule in order to enter a ‘slip state’^14^. In the latter, the motor re-engages with the same protofilament, but with the labeled and unlabeled heads in reverted order. However, as only one of the heads is tracked, the unpaired 8 nm step might also arise from kinesin constantly being attached to the same protofilament, but displaying an ‘inchworm’ step at this instant. A clearer case for the switching between protofilaments can nevertheless be made by the off-axis step (Fig. 2a,b (3) marked in green) at around 680 nm, which induces a shift of about 7 nm to the side, perpendicular to the assumed microtubule axis. This observation is best explained by kinesin-1 switching to another protofilament where it recovers its regular processivity^12,15^.

Pronounced protofilament switching is also displayed in other traces in Fig. 3a,b, where off-axis steps shift the straight stepping trajectories by up to 26 nm. As the rate of detected photons remains constant during the step, we can nearly exclude that the steps after the shift arise from a different motor protein. A possible interpretation is that the motor interrupts its movement, potentially due to a roadblock, detaches from the protofilament, and re-engages with the microtubule at another protofilament. With such pronounced off-axis displacements, it is likely that the motor fully detaches from the protofilament and diffuses along the microtubule surface before re-engaging– occasionally even on the opposite side of the microtubule (Fig. 3a). This rescue mechanism is of great interest as it indicates how kinesin-1 might regain processivity after premature detachment.

**Fig. 3.**
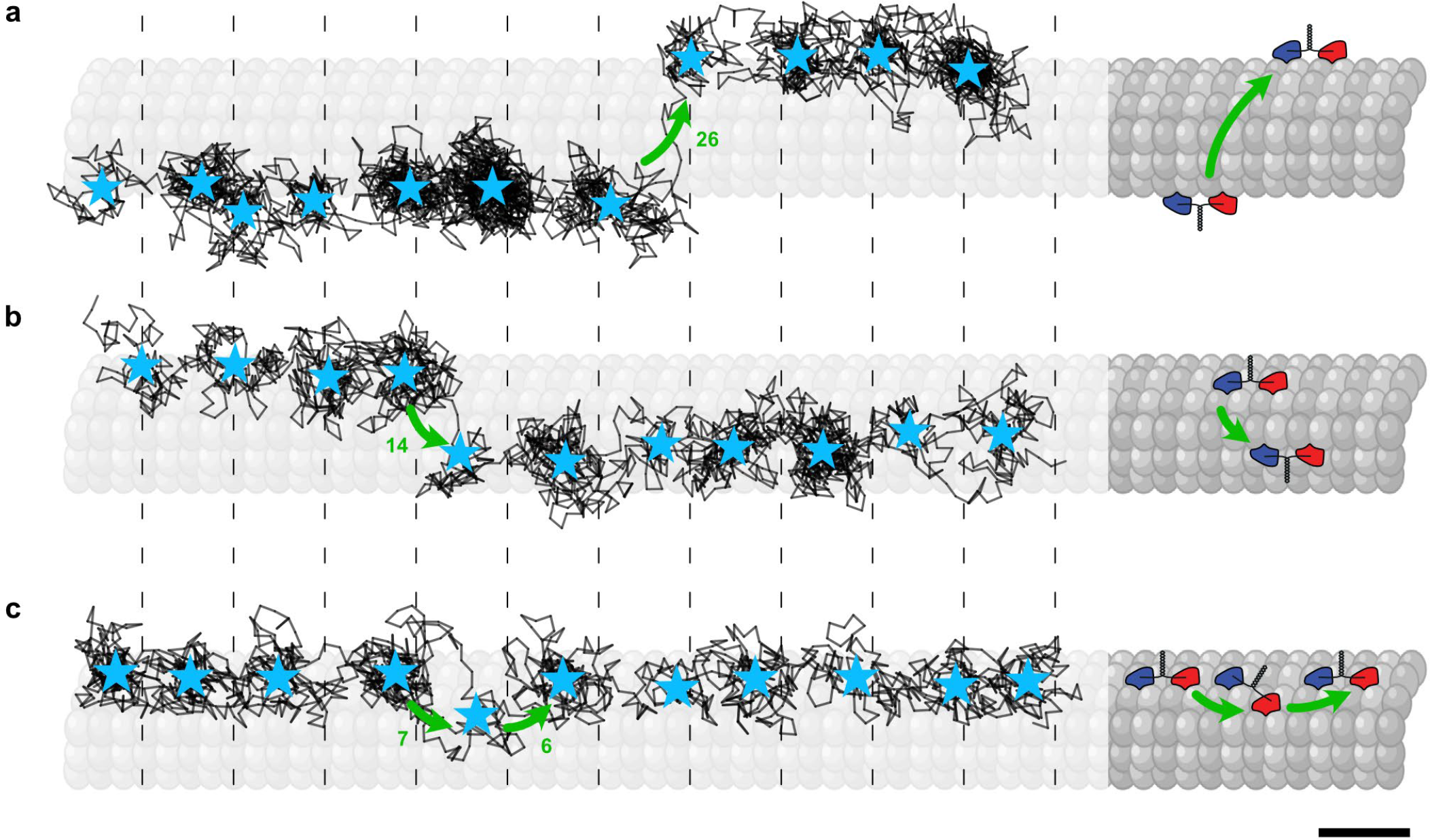
Observation of kinesin-1 off-axis movements. **a**, MINSTED localization trace revealing a 26 nm off-axis displacement on the microtubule (sketched in grey), while keeping its 16-nm stepping periodicity. Construct: E215C-ATTO 647N. **b**, Trace displaying a 14 nm off-axis step associated with an on-axis step of 8-nm (‘phase-shift’). Construct: E215C-ATTO 647N. **c**, Back and forth off-axis stepping with 7 nm and 6 nm steps, respectively. Construct: T324C-ATTO 647N. Individual localizations are shown as black line segments, whereas inter-step mean positions are shown as light blue stars. Schematics on the right of each panel are interpretations of each trace. Dashed lines mark 16 nm distances. Scale bar 16 nm.

Closer inspection of all recorded traces revealed that protofilament switching occurs about every 50 steps (15 protofilament switches out of 779 total steps). Sideward displacements occur both to the right and to the left, in accordance with earlier observations made from tracking of kinesin-1 labeled with gold beads or quantum dots^15,16^. Furthermore, for about half of the cases, the ∼ 16 nm periodicity was conserved, whereas in the other half, the subsequent pattern of binding sites was shifted by ∼ 8 nm after the switch. The apparent absence of a preferred binding site on the new protofilament supports the hypothesis that the motor is weakly bound to the microtubule when it switches protofilaments.

On rare occasions, kinesin-1 displayed a sidestepping where it seemed to fleetingly switch to another protofilament before going back to the initial one. For example, a significant off-axis displacement of about 7 nm was observed during a regular plateau after a 16 nm on-axis step (Fig. 3c). With the subsequent regular step, the off-axis displacement was reversed, apparently bringing the labeled head back to the previous protofilament. Based on this observation, we sketched a mechanism (Fig. 3c, right) suggesting how kinesin-1 can circumvent a (small) obstacle on a single protofilament.

Reasoning on the nature of the roadblocks, we note that our microtubules were polymerized from highly pure tubulin with < 1% of microtubule-associated protein (MAP) and a minimal fraction of biotinylated (10%) and fluorophore-conjugated (2%, Alexa Fluor 488) tubulin. Roadblocks may also arise from non-specifically bound proteins (BSA-bt, pyranose oxidase, catalase) or inactive and non-fluorescent kinesins. The minimally modified microtubule surface may also explain why the dwell time before a protofilament switch (median 20 ms) was not substantially increased (median dwell time of all steps 23 ms), unlike in a study where the roadblocks were permanent^15^. Offering insights on how the motor protein copes with roadblocks, our measurements suggest that the stepping of kinesin-1 is rather flexible.

Altogether, measuring nanometer steps with (sub)millisecond temporal resolution in traces ranging over a few hundred nanometers, MINSTED is highly suitable for studying (motor) protein dynamics and conformational changes. Moreover, our MINSTED tracking is rather unique in that the localization is updated for every detected photon, yet it delivers single-digit nanometer precisions in the raw trace. It remains to be studied whether the same performance can be accomplished with MINFLUX.

In any case, the deeply underlying reason why both MINSTED and MINFLUX localization require fewer detected photons is the same. Due to diffraction, defining a molecular coordinate with a diffracted light beam with high precision undeniably requires many photons. Whereas in conventional localization providing these photons is entirely up to the fluorophore, in MINFLUX and MINSTED the majority of the photons needed for localization is provided by the laser. Although this principle comes to full power in our MINSTED tracking, additional developments in fluorophore chemistry, the coordinate-finding algorithm, and the optical system are poised to improve the spatio-temporal resolution of coordinate-targeted fluorophore tracking even further.

## Author contributions

H. v. d. E., L. S., M. H., and M. W. performed the measurements. H. v. d. E. and M. W. built and operated the setups. L. S. prepared the kinesin samples and interpreted the motor protein results. H. v. d. E. and M. H. processed the raw data with feedback from M. W.. L. S. interpreted the data with feedback from S. W. H.. M. W. implemented the analytical model. S. W. H. initiated, outlined and supervised the research project. L. S., H. v. d. E., M. W., and S. W. H. wrote the manuscript. All authors contributed to the manuscript either through discussions or directly.

## Acknowledgement

We thank Miroslaw Tarnawski and the Protein Core Facility at the MPI Heidelberg for performing the QuikChange site-directed mutagenesis and protein expression. We also thank Sebastian Fabritz and the MS Core Facility at the MPI Heidelberg for recording the mass spectroscopy data and Antonio Politi from the Light Microscopy Facility at the MPI Göttingen for help with the TIRF recordings.

## Data availability

The data that supports Figs. 1-3 within this paper is available from the corresponding author upon reasonable request.

## Code availability

The custom simulation and analysis codes are available from the corresponding author upon reasonable request.

## Methods

### MINSTED microscopes

We used two previously described MINSTED fluorescence microscopes^7,11^ with each a different pair of excitation (λ = 560 nm/635 nm) and STED wavelength (λ = 636 nm/775 nm). Both microscopes featured a fast electro-optic 2D scanning system that positioned the co-aligned beam pairs with Angström precision in the focal plane. The lasers were pulsed with a repetition rate of 40 MHz/20 MHz repectively, while the excitation pulses and STED pulses had a duration of 0.1-0.2 ns and ≤ 1.5 ns, respectively. About 1 ns after each excitation pulse, a time gate for fluorescence detection with an avalanche photo diode was opened for 8 ns, to keep background low. In both setups the FPGA control modified the position of the co-aligned laser beam, as well as their power, in response to every detected photon. Lateral sample stability was ensured by tracking the position of metal colloids in the sample with back-scattered near-infrared light. A focus-lock, tracking the reflection of an NIR beam from the coverslip-sample interface, stabilized the sample along the optical axis. The sample was translated and actively position-corrected with sub-nm precision by means of a three-axis piezo stage.

### MINSTED measurements

MINSTED was implemented using the previously described localization algorithm^7^. All measurements were executed with a circle diameter 2*r* matching the FWHM of the employed E-PSF. The center positions were updated upon each photon detection with a stepsize corresponding to αα = 15% of the circling radius *r*. For the step temporal response measurements (Fig. 1d-f), the MINSTED localizations were initiated when four neighboring pixels in a confocal overview scan (1.6 ms dwelltime, 80 nm pixelsize) crossed an accumulated number of 40-80 counts. For kinesin measurements (Fig. 2,3) with Cy3B, the threshold was set to 4 (60 μs dwelltime, 80 nm pixelsize); while with ATTO 647N, it was chosen to 5 (60 μs dwelltime and 50 nm/75 nm x/y pixelsize).

### Sample preparation

#### DNA origami

10 μl of gold nanorods (A12-40-980, Nanopartz) at 0.2 mg/ml in methanol were dried onto a coverslip, that was previously cleaned in an ultrasonic bath with Hellmanex II (Hellma), and then treated with an air plasma for approx. 15 min. The coverslip was glued to a microscope slide using double-sided scotch tape, creating a flow channel. The latter was rinsed with PBS (1x PBS 7.4 pH) and then filled with 15 μl of 0.5 mg/ml biotinylated BSA (A8649, Sigma-Aldrich) in PBS. After 4 min incubation and washing with 100 μl PBS, the channel was filled with 15 μl 0.5 mg/ml streptavidin (11721666001, Sigma-Aldrich) in PBS, and incubated for another 4 min. The channel was flushed with 100 μl of 10 mM MgCl_2_ in PBS before incubating 15 μl of the DNA origamis (3×3 6 nm, GattaQuant) for 15 min. Thereafter, the channel was washed with 400 μl of 75 mM MgCl_2_ in PBS and the 15 μl of the imager strand solution was added. The latter consisted of 5 nM of Cy3B coupled to the 3’ end of a DNA oligonucleotide (P1 sequence: 5’-3’ CTAGATGTAT, Metabion) in an oxygen-deprived reducing-oxidizing buffer^17^. Each 200 μl of this buffer consisted of 100 μl reducing-oxidizing buffer (10% (w/v) glycose, 12.5% (v/v) glycerol, 0.1 mM TCEP, 1 mM ascorbic acid) and 100 μl PBS supplemented with 2 μl of oxygen removal enzyme mix (25 units of pyranose oxidase (P4234, Sigma-Aldrich) and 80 μl of catalase (C100, Sigma-Aldrich) with 170 μl of PBS), 1 μl of 200 mM methyl viologen dichloride hydrate (856177, Sigma-Aldrich) and 75 mM MgCl_2_.

### Kinesin

#### Expression of kinesin

Cysteine-light truncated (at amino acid position 560) human kinesin-1 constructs (hereafter referred to as kinesin) were expressed in E. coli using the plasmids K560CLM T324C (#24460, Addgene)^18^, K560CLM E215C (kindly provided by the Yildiz Lab, University of California, Berkeley), and K560CLM K28C produced from CLM RP HTR (#24430, Addgene)^19^ by QuikChange site-directed mutagenesis (using PfuUltra HF polymerase (600380-51), Agilent) as described by Tomishige & Vale^19^. All plasmids were verified by DNA sequencing. The constructs each contained a single solvent-exposed cysteine at amino acid position 324, 215 or 28 for labeling, and a C-terminal His_6_-Tag for purification (via 5 ml HisTrap FF (GE17-5255-01, Cytiva)). The purified protein (in 25 mM piperazine-N,N′-bis(2-ethanesulfonic acid) (PIPES, P-1851, Sigma-Aldrich) pH 6.8, 2 mM MgCl_2_ (1.05833.0250, Merck), 1 mM ethylene glycol-bis(2-aminoethylether)-N,N,N′,N′-tetraacetic acid (EGTA, E3889, Sigma-Aldrich), 0.1 mM ATP (BP413-25, Fisher Scientific), 0.2 mM TCEP (J60316, Alfa Aesar), around 300 mM NaCl (1.06404.1000, Sigma-Aldrich), 10% (m/v) sucrose (S1888, Sigma-Aldrich)) was aliquoted, flash frozen in liquid nitrogen and subsequently stored at -80°C.

#### Labeling of kinesin

Kinesin was labeled with ATTO 647N maleimide (AD 647N-41, ATTO-TEC) or Cyanine3B maleimide (19380, Lumiprobe) over night at 4°C. Excess dye was removed from the reaction mixture by size-exclusion chromatography (PD MiniTrap G-25, 28-9180-07, Cytiva) according to the manufacturer’s protocol. The degree of labeling (DOL) was determined by UV-Vis spectroscopy (DS-11+ Spectrophotometer, DeNovix) and mass spectrometry (ESI, maXis II ETD, Bruker). Sucrose was added to the labeled protein in a concentration of 10 % (w/v) and aliquots were flash-frozen in liquid nitrogen and stored at -80°C.

#### Preparation of microtubules

Biotinylated and fluorescently labeled microtubules were polymerized from 88% Cycled Tubulin (032005, PurSolutions, LLC), 10% Labeled Tubulin-Biotin-XX (033305, PurSolutions, LLC) and 2% Labeled Tubulin-Alexa Fluor 488 (048805, PurSolutions, LLC). The lyophilized tubulin variants were suspended in PEM80 buffer (80 mM PIPES, 0.5 mM EGTA, 2 mM MgCl_2_, pH 7.4) with 1 mM guanosine-5’-[(α,β)-methyleno]triphosphate (GMPCPP; NU-405S, Jena Bioscience) and the solution was incubated for 30 min at 37°C. Afterwards, the polymerized microtubules were centrifuged at 21,000x g in a bench-top microcentrifuge (Fresco 21, Thermo Scientific) for 15 min, washed with PEM80 and centrifuged at 21,000 × g for 15 min. The microtubule pellet was resuspended in PEM80, aliquoted, flash-frozen in liquid nitrogen and stored at -80°C.

#### Sample preparation

Flow chambers were constructed using oxygen-plasma-cleaned coverslips and double-sided adhesive tape. The chambers were incubated with 0.2 mg/ml biotinylated poly-L-lysine-polyethylene-glycol (PLL-PEG-bt) solution (PLL(20)-g[3.5]-PEG(2)/PEG(3.4)-biotin, Susos AG Inc.) supplemented with 1 % (v/v) Tween 20 (P9416, Sigma-Aldrich) in ddH_2_O for 15 min, rinsed with PEM80, incubated with 10 μg/ml neutravidin (NVD; 31000, Thermo Fisher) in PEM80 for 5 min, and rinsed with PEM80. The flow chambers were incubated with microtubules diluted in 20 μM cabazitaxel (FC19621, Biosynth Carbosynth) in PEM80 for 5 min, rinsed with PEM80, blocked with 100 μg/ml biotinylated bovine serum albumin (BSA-bt; A8549-10MG, Sigma-Aldrich) in PEM80 with 20 μM paclitaxel (10-2095, Focus Biomolecules) added for 30 min, and rinsed with PM15 buffer (15 mM PIPES, 2 mM MgCl2, pH 7.4).

Labeled kinesin in measuring buffer (1 mM 1,4-dithiothreitol (DTT, 6908.1, Carl Roth), 20 μM paclitaxel, 10 μg/ml BSA-bt, 1 mM methyl viologen, 1 mM ascorbic acid, 1 mM adenosine 5′-triphosphate (ATP; A3377-1G, Sigma-Aldrich) with an oxygen scavenger system (0.25 units pyranose oxidase (P4234, Sigma-Aldrich), 0.8 μl catalase from bovine liver (C100, Sigma-Aldrich) and 5% (w/v) D(+)-Glucose (HN06.1, Carl Roth)) in PM15 buffer) was added and the flow chamber was sealed with picodent silicone putty or nail polish.

## Data analysis

### Step temporal response (Fig. 1d-f)

The s = 16 nm dislocation of the co-aligned beams (mimicking a lateral fluorophore step) was controlled by the FPGA. This was accomplished by modulating the driving voltage of the x-axis electro-optic scanner every T = 15 ms. After cutting off all circle center positions where *r* had not yet converged to its final value, we discarded all the traces with a duration < 31 ms or containing < 100 detected photons. Around each remaining step, we segmented an interval of [-T, T] (if possible). The circle center positions along the x- and y-axis within each of those step intervals were denoted with X(t) and Y(t), respectively, with a detection time t relative to the time of the respective step. In addition to the original data with irregular detection times, the steps were mapped onto a regular, common time interval 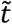with a sampling time of T/5000. The mapped positions 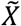 resulted from an interpolation algorithm, selecting the most recent spatial coordinate in *X* for each element in 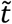. After segmentation, all steps were discarded having a detection rate < 70% of the median value among all localizations. Also, we discarded all steps featuring displacements of any 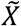 larger than (*s* + 2 std(*Y*)).

To adapt for the global position offset of each step, the mean value *x*_0_ = ⟨*X*(*t*)⟩_*Z*_ was calculated within the time interval Z = (−10 ms < *t* < 0), and subtracted from the respective step responses *X* and 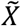 (i.e., zeroing). Artificially adding the step size s to all 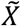 before the first photon arrival at all times 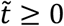 resulted in a set of both temporally and spatially overlaid steps, jumping from a position s down to a position 0 at 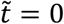. The mean step response was calculated as an ensemble average of the interpolated responses 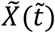. The decay time τ_0_ was found from the mean step response by linearly interpolating the time 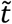 at which the mean position decayed to s/e.

With this initial estimate of the response time, the interval Z was adapted to (−*T* + 5τ_0_ < *t* < 0), and the zeroing was repeated in order to ensure convergence of the trace for all times within Z. The final response time τ was estimated (now from the corrected positional values) as above.

For the estimation of the single-photon spatial precision σ, the standard deviation of 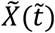 among the measured steps was evaluated. The resulting σ was computed as the mean of those standard deviations over 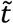 for 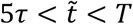. The step-localization spatial precision was computed as 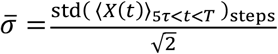. Assuming that all steps sample the same positional value (in the given time interval), the standard deviation over the temporal average would give the spatial precision of the mean position. Due to the statistical error in the zeroing step and as the uncertainty of the latter can be assumed equal to the step-localization spatial precision, a factor of 2^−1*/*2^ is multiplied.

#### Kinesin stepping (Fig. 2-3)

The raw tracks of the kinesin movement were cut by removing the first center positions where the sampling radius had not yet converged to its minimal value as well as the last 16 photons (in order to remove potential random motion due to pure background detection after a bleaching event occurred). Thereafter, multiple filters were applied, to identify traces that display actual kinesin movement with at least a few steps. First, all traces with a duration shorter than T_min_ were discarded. We performed an initial rotation operation on the remaining traces, by forcing the first and last center position to lie on the x-axis. We then estimated the covered distance from the minimal and maximal x deflection and filtered for minimal distances of D_min_. The single-photon spatial precision σ was approximated by the y-axis standard deviation. Traces with a single-photon spatial precision falling outside the interval [σ_min_,σ_max_], and traces with an aspect ratio std(X)/std(Y) larger than r_min_ were discarded. The current count rate k(t) was estimated as the reciprocal of the moving mean of each 20 values over the time differences of consecutive photon detection times and was further smoothed by applying a moving mean of 50. Another filter was applied to the standard deviation of k(t) in order to discard traces with large fluctuations 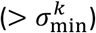 in brightness. Such traces are likely to be disturbed by the presence of multiple emitters in the sampling, or events where one emitter was lost and a second one found after a short dark period. The filtering parameters for the recorded datasets are presented in Table 1.

**Table 1:** Filtering parameters.

| Sample | $T_{\min}$ | $D_{\min}$ | $[\sigma_{\min}, \sigma_{\max}]$ | $r_{\min}$ | $\sigma_{\min}^k$ |
| --- | --- | --- | --- | --- | --- |
| E215C ATTO 647N | 50 ms | 64 nm | [2,10] nm | 4 | 2000 Hz |
| T324C ATTO 647N | 50 ms | 64 nm | [2,10] nm | 4 | 2000 Hz |
| K28C ATTO 647N | 50 ms | 64 nm | [2,10] nm | 4 | 2000 Hz |
| K28C Cy3B | 50 ms | 64 nm | [2,10] nm | 4 | 1800 Hz |

All remaining traces were fitted with a linear polynomial and rotated parallel to the x-axis. On the rotated traces, we performed an edge detection, using the function findchangepts (Matlab R2021b) in both x- and y-axis with the penalty parameter ‘MinThreshold’ set to 130 (⟨std(*Y*)⟩_t*r*aces_)^2^ and the ‘Statistics’ parameter set to ‘mean’. Detected steps in the y-axis were discarded if they referred to a detection index closer or equal to 20 photons with respect to an x-axis step. Finally, the mean positions between the detected steps were computed and those covering a Euclidian distance of ≤ 5 nm were discarded. Re-computing the mean coordinate between each two steps left us with an initial set of plateau positions. In order to retrace the local microtubule orientation for each single trace, we computed the angle of the stepping vectors between the plateau positions with respect to the x-axis and rotated each trace so that the median angle vanished. This enforced an alignment of the major stepping direction with the x-axis and mostly repressed an angular alignment with respect to off-axis stepping. With those final rotated traces, we repeated the edge detection described above.

